# Hierarchical Bayesian Integrated Model for Estimating Migratory Bird Harvest in Canada

**DOI:** 10.1101/2021.05.04.442620

**Authors:** Adam C. Smith, Thomas Villeneuve, Michel Gendron

## Abstract

The Canadian Wildlife Service (CWS) requires reliable estimates of the harvest of migratory game birds, including waterfowl, to effectively manage populations of these hunted species. The National Harvest Survey is an annual survey of hunters who purchase Canada’s mandatory migratory game bird hunting permit, integrating information from a survey of hunting activity with information from a separate survey of species composition in the harvest. We use these survey data to estimate the number of birds harvested for each species, as well as hunting activity metrics such as the number of active hunters and days spent hunting. The analytical methods used to generate these estimates have not changed since the survey was first designed in the early 1970s. Here we describe a new hierarchical Bayesian integrated model, which replaces the series of ratio estimators that comprised the old model. We are now using this new model to generate estimates for migratory bird harvests as of the 2019-2020 hunting season, and to generate updated estimates for all earlier years. The hierarchical Bayesian model uses over-dispersed Poisson distributions to model mean hunter activity and harvest (zero inflated Poisson and zero truncated Poisson, respectively). It also includes multinomial distributions to model some key components including, variation in total harvest across periods of the hunting season, the species composition of the harvest within each of those periods, and the age and sex composition in the harvests of a given species. We estimated the parameters of the Poisson and the multinomial distributions for each year as random effects using first-difference time-series. This time-series component allows the model to share information across years and reduces the sensitivity of the estimates to annual sampling noise. The new model estimates are generally very similar to those from the old model, particularly for the species that occur most commonly in the harvest, and so the results do not suggest any major changes to harvest management decisions and regulations. However, estimates for all species from the new model are more precise and less susceptible to annual sampling error, particularly for species that occur less commonly in the harvest (e.g., sea ducks and other species of conservation concern). This new model, with its hierarchical Bayesian framework, will also facilitate future improvements and elaborations, allowing the incorporation of prior information from the rich literature and knowledge in game bird management and biology.

---

Reliable estimates of the harvest of migratory bird populations are necessary for Canada to manage populations of migratory game birds and to meet its commitments under the Migratory Bird Convention Act (Migratory Birds Convention Act, 1994, S.C. 1994, c. 22). Declining waterfowl populations in the early 1900’s was the first indication that there could be negative effects on the sustainability of some migratory bird populations if there was no protection against excessive hunting (Nichols et al. 1995, Cooch et al. 2014). Under the Migratory Bird Convention Act, the Canadian Wildlife Service (CWS) has implemented regulations in order to prevent collapses of such populations while simultaneously allowing for recreational hunting. Management actions include designated hunting seasons, daily bag limits and possession limits.

Harvest data and estimates of harvest are increasingly being used as one of the main data sources for estimating various parameters of population dynamics, such as determining population size (Alisauskas et al. 2009, 2014, Zimmerman et al. 2017), fecundity (Zimmerman et al. 2010, Osnas et al, 2016), sex ratio (Hagen et al. 2018), as well as in integrated population modeling efforts (Saunders et al. 2019). Reliable estimates of harvest are particularly important for harvested species with no reliable, population-wide count-based surveys, such as many Arctic-breeding geese.

The CWS estimates the annual recreational harvest of migratory game birds in Canada using the National Harvest Survey (NHS). The program uses information gathered from purchasers of the Migratory Game Bird Hunting Permits (MGBHP). These permits are mandatory for all non-indigenous individuals to hunt migratory game birds in Canada, and so they provide a sampling universe from which the survey can draw (Sen 1976, Cooch et al. 1978). The NHS was initiated in 1967, and after several years of fine-tuning, has been conducted annually using the same analytical methods since 1976 (Sen et al. 1975). The National Harvest Survey website (https://wildlife-species.canada.ca/harvest-survey) is the central platform that annually distributes a variety of estimates related to harvests and hunting activity. For example, published estimates include: the number of active waterfowl hunters, the number of waterfowl hunting days, the number of successful hunters, species-specific harvest, and age-ratios. Providing reliable estimates is a critical part of the National Harvest Survey program since they, along with other CWS monitoring programs, are used to assess the status of migratory game bird populations in Canada (Canadian Wildlife Service Waterfowl Committee. 2020) and levels of sustainable harvest (Gilliland et al. 2009, Palumbo et al. 2020).

The analytical methods used to generate estimates of migratory game bird harvest could benefit from contemporary model-based approaches such as hierarchical Bayesian models (Dorazio et al. 2016). These model-based hierarchical approaches provide a coherent framework for sharing information through time, and among geographic strata and/or hunter classes (Cressie et al. 2009). Bayesian approaches provide both improved estimates of uncertainty and a transparent and explicit way to incorporate the ecological and sociological knowledge (Gelman et al. 2013, van de Schoot et al. 2021) that comes from the rich history of game bird population biology and harvest management in North American (Nichols et al. 1995, NAWMP 2018). Since the beginning of the survey, the estimates of harvests and hunting activity have been made using design-based equations for ratio estimates (Cochran 1977), which we refer to here as the “old model”. The relevant ratios included the ratios of harvest per hunter, hunting-days per hunter, and the number parts of a given species per total number of parts (Sen et al. 1975). This design-based old model calculated these mean ratios for each stratum of a stratified random sample in a given year (Cooch et al. 1978). However, each year’s estimates were derived independently of all other years, and were therefore particularly sensitive to variation among years in the sample, response rates to the survey, and the total number of hunters.

In this study, we describe a new hierarchical Bayesian integrated model (hereafter “the model” or “the new model”) that estimates annual harvest of all species of migratory game birds hunted in Canada, as well as summary estimates of hunter activity and total number of ducks, geese, and other major species groups. With this new model, we expect to generate estimates that are more precise and less sensitive to sampling noise, particularly for species of conservation concern or those less abundant in the harvest. We provide the full code required to run it in an online supplement, which greatly increases the transparency of these estimates over the old model, which was never published in a formal and comprehensive way. In addition, the Bayesian framework of this new model will allow for future improvements using informative priors that incorporate the ecological and sociological knowledge that underlies the long history of waterfowl harvest management in North America (NAWMP 2018).

## METHODS

### Overview of the NHS

The NHS sampling methods have remained the same since 1976. It is separated into two primary components: the harvest questionnaire survey (HQS), and the species composition survey (SCS). Hunters responding to the HQS provide information on the total harvest of broad groups of species (e.g., all ducks, geese, and other non-waterfowl species), where they hunted, and the total number of days spent hunting, as well as calendar information indicating how many birds in each group they harvested on each day of the season. We stratified the HQS responses based on hunter residency (Canadian vs US residents) and their previous hunting activity. Hunters responding to the SCS provide information on the species composition of their harvest by submitting wings (ducks and murres) or tail-fans (geese) that are identified to species, and if possible, aged and sexed, by waterfowl biologists. We then integrated the overall hunting information of the HQS with the species composition information of the SCS to generate estimates of the species-level harvest in each year.

We randomly selected potential participants for the HQS and the SCS using records in the MGBHP database. We separated permit holders into 24 geographic hunting zones (Fig. 1) and for the HQS, we further separated them into four classes (Table 1). We selected hunters in categories A and E from the current year’s permit records, while those in categories B and D are selected from the previous year’s permit records. This classification is based on their country of residence and whether or not they held a permit in previous years. The precision of the estimates increase by grouping the hunters into one of the four classes because it takes into account differences in hunting activity and success.

**Figure 1:**
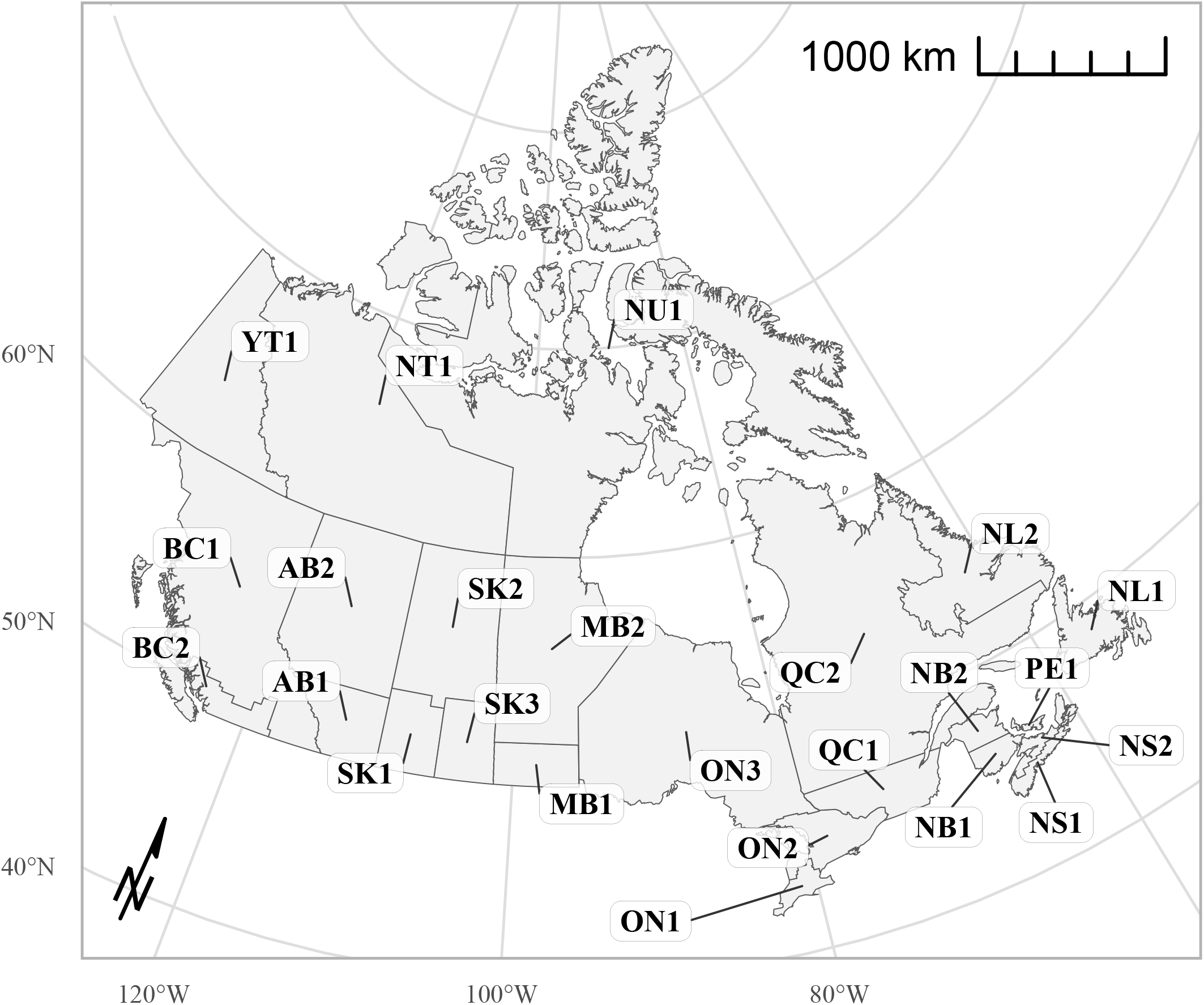
Map of the 24 geographic hunting zones in Canada. The zones represent divisions of Canada’s provinces and territories. The provinces and territories are labeled as follows: AB = Alberta, BC = British Columbia, MB = Manitoba, NB = New Brunswick, NS = Nova Scotia, NT = Northwest Territories, NU = Nunavut ^a^, ON = Ontario, PE = Prince Edward Island, QC = Quebec, SK = Saskatchewan, and YT = Yukon. ^a^ there are insufficient data to fit the model in Nunavut

**Table 1:** Hunter class description for the harvest questionnaire survey.

| Sample code | Description of hunter class | Sample drawn from |
| --- | --- | --- |
| **A** | Residents who did not purchase a permit in the previous year | Current year |
| **B** | Residents who bought a permit year previously but not in the year prior to that | Previous year |
| **D** | Residents who bought a permit in the preceding 2 years | Previous year |
| **E** | Non-residents | Current year |

We selected SCS participants differently than those in the HQS. In order to distribute plastic envelopes for the wing and/or tail samples before the start of the hunting season, we selected participants from the previous year’s MGBHP database. First, we determined a hunter’s willingness to participate in the survey by mailing a participation screening card in early July. We generated this random sample of permit holders based on survey participation history, hunting success, and permit renewal status (Table 2). We only sampled Canadian residents in the SCS because of the challenges in delivering envelopes and receiving bird-parts across international borders. Hunter selection is biased toward hunters who previously cooperated. This is beneficial because it increases the response rate, the estimate precision, and the cost efficiency of the survey. However, to limit the bias caused by repeatedly re-sampling the same hunters, we removed two-year SCS participants from the sample for at least one year.

**Table 2:** Hunter categories for species composition survey.

| **Sample** | **Description** |
| --- | --- |
| **SA** | SC, SD, SE, or SF respondent in the previous year |
| **SC** | HQS respondent in the previous year who shot more than five waterfowl |
| **SD** | HQS respondent in the previous year who shot one to five waterfowl |
| **SE** | Renewal hunter of the previous year, not eligible for SA, SC, or SD |
| **SF** | Non-renewal hunter of the previous year not eligible for SA, SC, or SD |

Due to changes over time in how permits have been purchased and hunters have been sampled for the survey, the allocation of permitted hunters and survey responses to each zone has also varied through time. Until recently, we sampled hunters based on the province and zone where they purchased their permit, because no information on the location of hunting activity was available at the time of sampling. Hunters who were sampled for the HQS indicated where they did most of their hunting. In general, the zone of purchase was predominantly the same as the zone of harvest. In recent years, many permits have been sold through an online portal where the zone of purchase is not relevant, and so we started asking hunters to indicate at the time of purchase where most of their hunting activity will take place. We then used this information to link a permit record to a sampling zone. In the new model, we included transformation factors to account for the proportion of hunters that were sampled in a different zone than the one in which they hunted. These transformation factors will become less relevant once the transition to online-sold permits is completed and we can link all permits records to hunting zones based on intended location of hunt.

### The New Model

Using the new hierarchical Bayesian integrated model described here, we estimated the mean group-level (e.g., all ducks) harvest of hunters using an over-dispersed, zero-inflated, Poisson distribution, and the data from the same hunter on total days (days spent hunting) as an over-dispersed, zero-truncated Poisson distribution. We used the HQS calendar responses (how many birds were harvested on each day of the season) to estimate the proportions of the total harvest that occurred within each of a series of discrete periods of the season using multinomial distributions. We then portioned this period-specific total harvest across species by integrating with the SCS data on the number of birds of each species harvested within each period using a series of additional multinomial distributions. We corrected for the period-specific portioning because of the known decline in response rates to the SCS over the course of the season. Finally, we summed the period-specific total harvest estimates across periods to estimate the annual harvest of each species. We included in each component of the model (e.g., total harvest, total days, period-proportions of the total harvest, species proportions in a given period) an explicit random-walk time-series sub-model that shares information between sequential years. This time-series component assumes that each of these parameters has some relationship to the same parameter in the previous year, owing to long-term patterns in species abundance, hunter behaviour, and hunting regulations. For example, the estimated mean number of days spent hunting in a given year is a function of the same value in the previous year, plus some random error. These random-effect time-series components allow the model to share some information through time, while still allowing for any shape of year-to-year change, including smooth trends (e.g., declines in overall hunting through time), annual fluctuations (e.g., seasons with very poor weather that may have reduced activity or harvest success), and step-changes (e.g., introduction of new regulations that change harvest). Finally, we transformed the per-hunter mean values for group-level and species-level harvest to population-level estimates of total harvest using information on the total size of the permit-population hunting in a given zone (i.e., the total number of permitted hunters hunting in a given zone). We apply the model separately for the province and zone where harvest-activity took place.

To simplify the following sections in which we detail the specific components of the model, we use ducks as the example group. However, the model is the same for the other two groups of species, geese and murres (in Newfoundland and Labrador Zone-1), that include calendar and species-composition information.

### Total Harvest and Total Days

Each HQS response includes data on the total reported number of ducks harvested by hunter - *h*, in class – *c*, and year – *y*, which we modeled as an over-dispersed, zero-inflated, Poisson distribution:

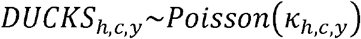

We modeled the mean of the Poisson distribution with a log-link, as a function of the year-effect (*α*_*y*_), an annual class-effect (*β*_*c,y*_), and an observation-level hunter-effect (*χ* _*h,c,y*,_), plus an offset for the log of the estimated mean number of days reported by the same hunter (*log*(*λ*_*h,c,y*_)), and a zero-inflation component (*z*_*h,c,y*_), which is the outcome of a Bernoulli trial and equals 1 with estimated probability of 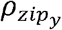 in year -*y*. The value 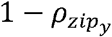 represents the proportion of active hunters that harvest no ducks in that year, in addition to the zero-harvests expected from the Poisson distribution. This extra parameter modeling zero-harvests accounts for active waterfowl hunters that only hunt one group of waterfowl (e.g., only hunt geese and not ducks), and must be modeled because the survey does not ask hunters to indicate what types of waterfowl they prefer to hunt: geese, ducks, or both.

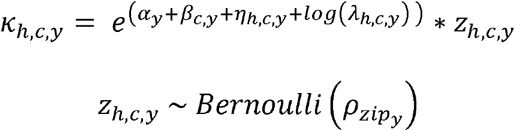

Each HQS response also includes data on the total reported number of days spent hunting by hunter - *h*, in class – *c*, and year – *y*. We modeled these data on number of days hunting as an over-dispersed, zero-truncated, Poisson distribution:

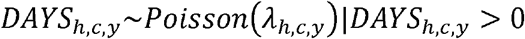

We modeled the mean of the zero-truncated Poisson (-*λ*_*h,c,y*_) with a log-link, as a function of the year-effect (*γ*_*y*_), an annual class-effect (*δ*_*c,y*_), and an observation-level hunter-effect (*ε*_*h,c,y*_).

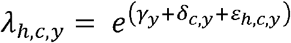

We modeled the year-effects for both harvest and days using a random-walk, first-difference time-series sub-model. We estimated the year-effects in the first year (*α*_1_ and *γ*_1_) as fixed-effects with a zero-mean, normally-distributed prior with a variance of 10 (e.g., *α*_1_∼*N*(0,10)). We estimated the remaining year-effects in year-y as random effects with a mean equal to the year-effect in the previous year and an estimated variance (e.g., 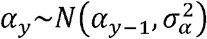). We set the priors for the variance of the year-effects as weakly informative priors on the standard deviations, following Gelman et al. (2006), using a half t-distribution with mean = 0, variance = 0.5 and degrees of freedom = 50 (*σ*_*α*_∼|*t*(0,0.5,50)|). This prior places approximately 95% of the prior density at values < 1.0, but includes a relatively long tail that allows for much larger values, if supported by the data. Given the common scale of parameter estimates in a log-link model such as this one, standard deviation values > 1.0 for random effects are extremely unlikely and so this prior is only very weakly informative (Gelman et al. 2006). We used this weakly informative prior on all sigma values in the model (e.g., *σ*_*δ*_, *σ*_*β*_, and *σ*_*ε*_ below).

We fixed the class-effect parameters contributing to the days and harvest components for the class with the largest number of hunters (class-D) at 0 in all years, so that the remaining class-effects were estimated as departures from the largest class (*β*_1,*y*_ =0 and *δ*_1,*y*_ = 0). We modeled the class-effects for the remaining classes in each year as normally distributed random-effects with a class-specific hyperparameter mean and estimated variance 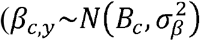 and 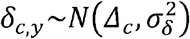. We gave the hyperparameter means (*B*_*c*_ and *Δ*_*c*_) normally-distributed priors with a variance of 10 (e.g., *B*_*c*_ ∼*N*(0,10)).

We estimated the observation-level hunter-effect parameters for days and harvest as zero-mean, t-distributed random-effects with class-specific variances and degrees of freedom 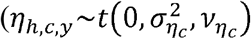 and 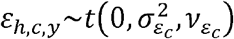). Using the t-distribution to model these over-dispersion effects allows the modeled hunter-level variation to fit heavier tails than a normal distribution (i.e., a greater number of extreme values in the tails of the distribution than predicted by a normal distribution). We used a normal distribution in early versions of the model, but in most zones and years the empirical distributions of these hunter-level effects showed much heavier tails than a normal distribution. These heavy tails capture the influence of particularly active and successful hunters, in particular. We gave priors to the degrees of freedom parameters (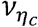 and 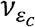) with gamma distributions with shape and scale set to 2 and 0.2 respectively. We estimated the parameters of the zero-inflation for the harvest 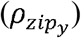 using a logistic regression sub-model that used a random walk time-series to track changes in these parameters over time. We estimated the logit of the first year zero-inflation parameter 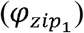 as a fixed effect in year-1 with a half-Cauchy prior following Gelman et al. (2008), which is a weakly informative prior with a reasonable scale for logistic regression coefficients. In all subsequent years, the logit of 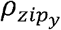 was a function of the value in the previous year, plus random variation.

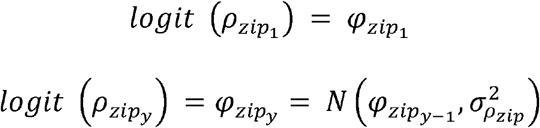

### Correction Factors to Estimate Number of Active and Successful Hunters

We estimated the binomial probability that a hunter in class-c and year-y actively hunted 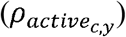 using the known numbers of HQS respondents who purchased a permit in that year, and the number of respondents who indicated they had > 0 days of hunting 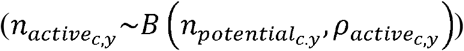. For each class-c, we modeled the series of 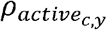 values for each year using a logistic regression sub-model similar to the one used for the zero-inflation component.

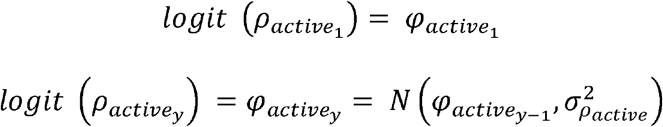

In the same way, we estimated the binomial probability that a hunter in class-c and year-y who actively hunted was successful (i.e., harvested > 0 birds, 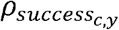) using the known numbers of HQS respondents who indicated they had > 0 days of hunting, and the number of HQS respondents who indicated they harvested > 0 birds 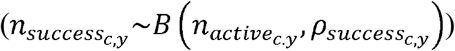. For each class-c, we modeled the series of 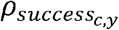 values for each year using a logistic regression sub-model identical to the one used for the proportion that were active.

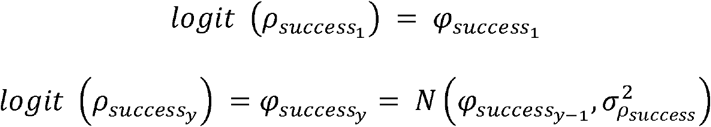

Similarly, we used similar time-series, logistic regression models to correct for inter-zone hunting. We modeled the annual probability that hunters sampled in a given zone would hunt mostly outside that zone 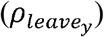 and the annual probability that a hunter hunting mostly inside the zone was sampled outside that zone 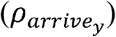. We derived the data for these two sub-models from a cross tabulation of the zone of hunt and the zone of sampling for all respondents. So for a given zone and year, we modeled the probability that a hunter sampled in that zone would hunt outside the zone 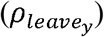 as a binomial distribution using the number of HQS respondents that were sampled in the zone 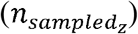 and the number of HQS respondents 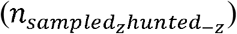 sampled in the zone who hunted outside the zone 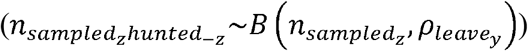. We also modeled the annual values of 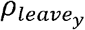 and 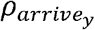 using a time-series logistic model, with a structure identical to the one used for the other binomial probabilities.

We combined the correction factors for inter-zone hunting, the proportion of active hunters, and the known total number of permits purchased in each year (*N*_*c,y*_) to estimate the total number of active hunters in a given class and year (*A*_*c,y*_).

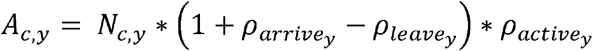

We re-scaled all mean values for hunters, classes, and species to population-level totals using the estimated number of active hunters for a given class and year (*A*_*c,y*_).

### Derived Totals of All Group-level Harvest and Activity

We calculated the estimated mean number of days hunting waterfowl (*<u>d</u>*_*c,y*_) for class-c and year-y as a derived statistic using the exponentiated sums of the relevant parameters, plus some added variance components to account for the asymmetries in the retransformation.

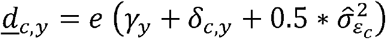

Where the term 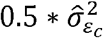 is an approximation of the half-variance retransformation from the mean of a log-normal distribution to the mean of the normal, accounting for the t-distributed overdispersion term in the model (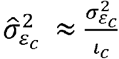, where 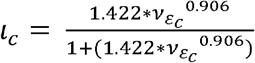). This term is an approximation because the variance of a t-distribution is undefined for some values of the degrees of freedom (Link et al. 2020). We used the same approximation suggested by Link et al. 2020, which they derived by estimating the parameters of a fitted regression model. Although this re-transformation is an area of ongoing research, we have found that it generates estimates of total harvest and activity that are comparable to estimates from the old model.

Similarly, we calculated the estimated mean number of birds (e.g., number of waterfowl) harvested by hunters in class-c and year-y (*<u>k</u>*_*c,y*_) using both the variance component related to days 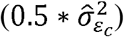 and a similar variance component for harvest 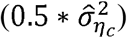. In this case, we also included the estimated probability of a non-zero harvest to account for the zero-inflation.

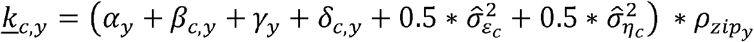

### Harvest by Species and by Age and Sex

We divide the estimates of mean total harvest by year and class into species-specific estimates of harvest using data on the seasonal harvest patterns from the HQS calendars (daily estimates of the number of waterfowl harvested for each respondent) and data on the species-composition information collected from the SCS. On the SCS envelopes in which wing and tail fans are received, hunters also provide information on their current year permit numbers as well as the date and location of harvest. These parts are then identified to species, and in most cases, grouped into demographic categories of age and sex by CWS biologists (Carney 1992). The SCS provides more reliable information on the species composition of the harvest in each zone and at different times of year than would be provided by asking hunters what species they harvested (Smith et al. 1974, Ahlers and Miller 2019). Not all parts can be identified to their demographic categories. We used any available information from each part, so for example the total number of parts summed across all demographic groups within a species will not necessarily equal the total number of parts for that species. Similarly, not all surveys complete the calendar portion of the survey that allows us to estimate the distribution of harvest across the season, but those survey responses can still be used to estimate the total harvest and total days. Our approach that uses all available information assumes that missing components (e.g., calendar information or age information for males and females) are missing at random. This assumption is also used in the old model, and it simplifies the new model and allows us to maximize our sample sizes in estimating the various proportions, such as the harvest in each period or sex ratios.

For calculating the species composition, we divided the harvest season into periods to account for the declining response rates to the SCS as the season progresses (e.g., due to response-fatigue, depleted initial envelope supply, etc. Smith et al 1975). The periods are important to account for differences in the phenology of migration and harvest among species, especially in zones where the hunting season is prolonged. The model generates unbiased estimates of the overall species composition across the entire season by combining separate estimates of the species composition in each period with unbiased estimates of the total harvest in these same periods (Cooch et al. 1978). Without this seasonal correction, the harvest of early migrants that are primarily harvested early in the hunting season such as, blue-winged teal (*Spatula discors*) and wood duck (*Aix sponsa*), would be over-estimated while the harvest of species that migrate and are harvested later in the season, such as scaup (*Aythya spp*.) and common eider (*Somateria mollissima*), would be under-estimated. We based the periods for a given zone and harvest group (e.g. ducks) on weekly divisions that included at least 5% of the total submitted parts. To allow the sharing of information on the seasonal patterns in harvest across years, we kept the periods consistent across all years. For ducks, we divided the season into 6 to 13 periods, depending on the length of the season in a given zone, and the distribution of the cumulative harvest across the season. In most cases, the earlier periods in a zone each represented a single week and the later periods may include more than 1 week. The final period often included many weeks, to capture the low level of hunting that continues through the winter.

The model includes three similarly structured sub-models that rely on multinomial distributions to estimate three key proportional distributions: 1) the proportions of annual harvest that occurred in each period using the calendar data; 2) the proportion of the harvest in each period that can be attributed to each species; and 3) the proportion of each species harvest that can be attributed to the age and sex categories (e.g., adult-female, immature male, etc.). We then combined these proportions with the total harvest estimates to estimate the number of birds harvested for each species in each year, as well as the number of birds in each demographic category for a given species.

For the sub-model for the proportional distribution of the harvest across W-periods, we used data on the number of harvested birds reported in each period-w, by hunter-h in year-y (*DUCKS*_*w,h,y*_), and the total yearly number of harvest birds reported by the same hunter (*DUCKS*_*h,y*_).

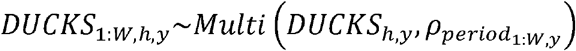

We gave Dirichlet-priors to the individual probabilities for each period-w and year-y 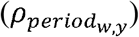, and we estimated the parameters of the Dirichlet using a log-link, random-walk time-series sub-model, similar to the parameters of the logistic regression for the binomial probabilities.

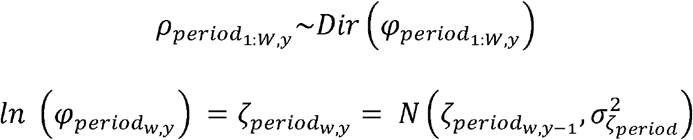

In the first year of the time-series, we kept the value for 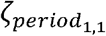 fixed at 0 for the first period and estimated as a fixed-effect for all other periods using a normal prior with mean of 0 and a variance of 10 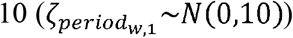.

For the sub-model for the proportional distribution of all S-species in period-w and year-y, we used data on the number of submitted parts of each species-s, period-w, in year-y (*PARTS*_*s,w,y*_), and the total yearly number of harvest birds submitted in that period (*PARTS* _*w,y*_).

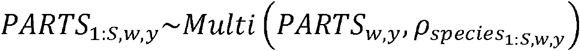

We gave Dirichlet-priors to the individual probabilities for each species-s in period-w and year-y 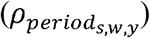.We estimated the parameters of the Dirichlet using a hierarchical, log-link, random-walk time-series sub-model, similar to the sub-model for the period distributions. However, in this case, we included a time-series year-effect (*τ*_*s,w,y*_) as well as a random effect for the mean abundance across periods for a given species (*ϕ*_*s, w*_).

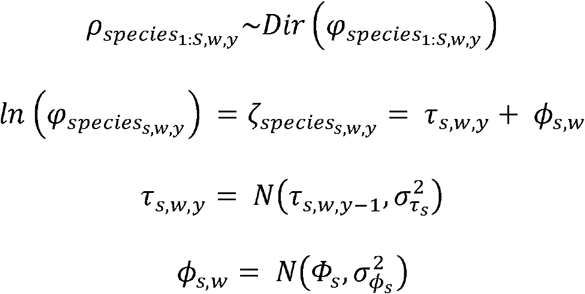

In the first year of the time-series, we estimated the value for the year-effect component in each period and species as a random effect using a prior with mean = 0 and a variance specific to the period 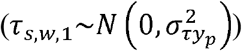. For the species mean abundance component (*Φ*_*s*_), which is the hyperprior for the random species effect by period, we fixed it at 0 for the first species, and we estimated it as a fixed effect for all other species, using a normal prior with mean of 0 and a variance of 10 (*Φ*_*s*_ ∼*N*(0,10)).

Finally, we gave Dirichlet-priors to the individual probabilities for each demographic group-d (i.e., each combination of the age and sex categories, such as adult-female, immature-male, etc.) and year-y 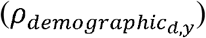, and we estimated the parameters of the Dirichlet using a log-link, random-walk time-series sub-model, which is identical to the parameters for the distribution of harvest across the periods.

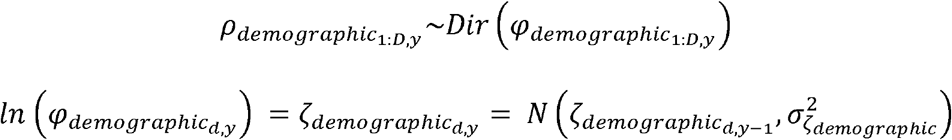

In the first year, we set the value for 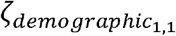 fixed at 0 for the first period and we estimated as a fixed-effect for all other demographic groups using a normal prior with mean of 0 and a variance of 10 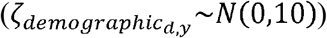.

### Final Estimates of Harvest and Activity

Finally, for the derived estimates, we re-scaled the relevant mean values (e.g., mean harvest of ducks by class-c in year-y, *<u>k</u>*_*c,y*_) to a total estimated harvest of ducks by class-c in year-y (*K*_*c,y*_) using the estimated number of active hunters in the same class and year (e.g., *K*_*c,y*_ = *<u>k</u>*_*c,y*_, ∗ *A*_*c,y*_). We summed these derived estimates of harvest, days, number of successful hunters, etc. across classes, zones, and provinces to generate the full suite of estimates provided to the public in the annual analysis.

Uncertainties for each of the estimates represent summaries of the full posterior distributions. Since we estimated the zone-level analyses independently using the same MCMC process, we summed the zone-level estimates for each posterior-draw of the MCMC, to estimate the full posterior distributions of the provincial and national estimates.

We implemented the MCMC analysis in JAGS (Plummer 2003), run through R (4.0.2), using the package jagsUI (Kellner 2019). We used a burn-in of 5000 iterations, and retained 3000 posterior-draws from 3 independent chains, thinned at a rate of 1/10. We assessed convergence within and across chains by visualizing trace plots and ensuring Rhat statistics for all interpreted parameters were < 1.1 (Gelman and Rubin 1992). We archived all anonymized survey data and code required to run the analyses reported here in a public repository (https://zenodo.org/badge/latestdoi/254251374).

### Model Checking and Model Fit

To assess the fit of the new model to the data, we employed posterior predictive checks within each of the main sub-models (Conn et al. 2018). For example, in developing the sub-model for the total number of ducks harvested, posterior predictions from earlier fitted models that lacked the zero-inflation component generated too few zero-valued harvests, compared to the observed data. Similarly, earlier versions of the new model that used a normal distribution to model the hunter-level over-dispersion effects generated predicted distributions of hunter level effects that did not fit the observed distribution of reported harvests well. In this case, the observed data tended to have many more high harvest estimates than predicted from the normal distribution. As a result, we adjusted the model to use a t-distribution with heavier tails than the normal to allow the model to predict the relatively large number of high harvests. Comprehensive assessments of fit for models that integrate multiple sources of data are complicated, for example any cross-validation approach requires a decision of what constitutes a unit of the data (Zipkin et al. 2021). In our case, there are nine independent response variables that the model predicts (number of birds harvested, number of days spent hunting, number of birds harvested each day of hunting, number of parts for each species, period, and year, number of parts identified to each demographic group within each species, plus four relatively simple counts of the numbers of hunters used to estimate the various binomial probabilities, such as 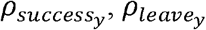, etc.). We anticipate some interesting comparisons of prediction error between this model and future modifications, but given the fundamental differences in modeling frameworks between the old and new models, we have focused on comparisons of the estimates and not predictions of data.

### Other Species

As part of the HQS, we also collect information on the harvest of other species of migratory birds, including Wilson’s snipe, American woodcock, sandhill crane, mourning dove, American coot, rails, and band-tailed pigeon. We capture this harvest information using a simplified portion of the questionnaire that collects information on the combined harvest effort for all non-waterfowl species and the harvests of each of the above species. In order to estimate the hunting effort and harvest for each of the non-waterfowl species, we used a simplified version of the new model. In this simplified version, we removed the species composition and calendar components, collapsed some of the hunter strata when samples sizes were small, and included an additional data matrix to ensure zero-harvest estimates in years when there was no allowable harvest for a particular species (i.e., a closed season in a particular zone and year). Otherwise, the components of the simplified new model that estimate harvest and activity through time are the same as they are in the new model for waterfowl and murres. We also included the complete code to run this simplified version of the new model in the online repository.

### Complete Estimates for 1976-2019

We have applied the new model to all data from 1976 – 2019 and generated estimates for all species and group-level harvests, all activity measures (e.g., total hunting days), and all age and sex ratios. We archived this full set of updated estimates, and the code to generate them, online (https://zenodo.org/badge/latestdoi/340784801). Annual updates of these estimates will be made available through the Canadian Wildlife Services harvest survey results website (https://wildlife-species.canada.ca/harvest-survey).

## RESULTS

The estimates of harvest derived from this new hierarchical Bayesian integrated model are generally similar to those from the old model, for data-rich species, zones, and years. For example, annual estimates of national harvests for total ducks and total geese are very similar across the entire time-series (Fig. 2), and over the last decade (2010-2019) the range of the annual percent-difference between new and old models includes both positive and negative values for all duck species (Fig. 3). For the species that are harvested commonly in large numbers, the differences between the two models are relatively small (i.e., species near the top of Figure 3). On average over the last decade (2010-2019), annual estimates of mallard (*Anas platyrhynchos*) and American black duck (*Anas rubripes*) harvest at a national level are 3-5% lower with the new model compared to the old model, and estimates for most other species are generally higher (i.e., species near the bottom of Fig. 3). The new model generally reduces the harvest estimates of the most commonly harvested species in a given zone (e.g., mallard or American black duck in most zones) by a small amount, and redistributes that harvest to species that may have been missing from the old model estimates in some years (Fig. 3). The new model also tends to reduce the magnitude of the annual fluctuations for species and zones that are relatively data-sparse, in comparison to the old model (e.g., northern pintail [*Anas acuta*] in Saskatchewan Zone 3; Fig. 4), but the sharing of information through time in the new model has less influence on the estimated annual fluctuations for the most commonly harvested species (e.g., mallard harvest in Saskatchewan Zone 3 and small-bodied Canada goose harvest in Manitoba Zone 1; Fig. 4). Additionally in cases with extremely sparse data, the new model generates non-zero estimates for species that happen to be absent from the species composition survey in a given year (e.g., black scoter [*Melanitta americana*] in Newfoundland and Labrador Zone 2; Fig. 4). The reduced annual fluctuations are particularly apparent for some species of conservation concern (Fig. 5).

**Figure 2.**
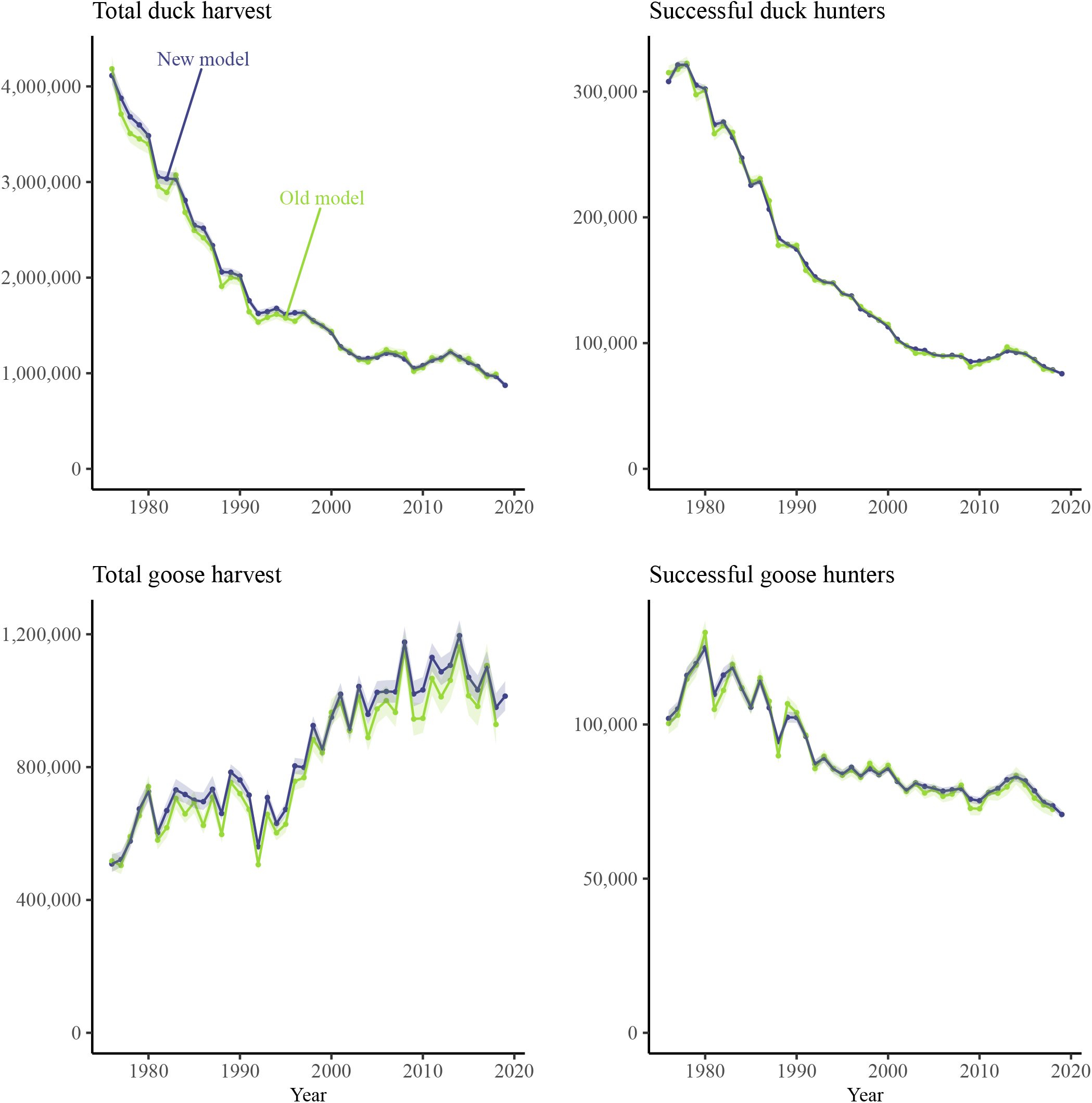
National estimates of the total harvest and total number of successful hunters for all duck species and all goose species from 1976-2019 from the Canadian National Harvest Survey, using the new hierarchical Bayesian model described here (darker line) and the old model that it replaces (lighter line). The semi-transparent ribbon surrounding each line represents the 95% credible/confidence intervals on the estimates.

**Figure 3.**
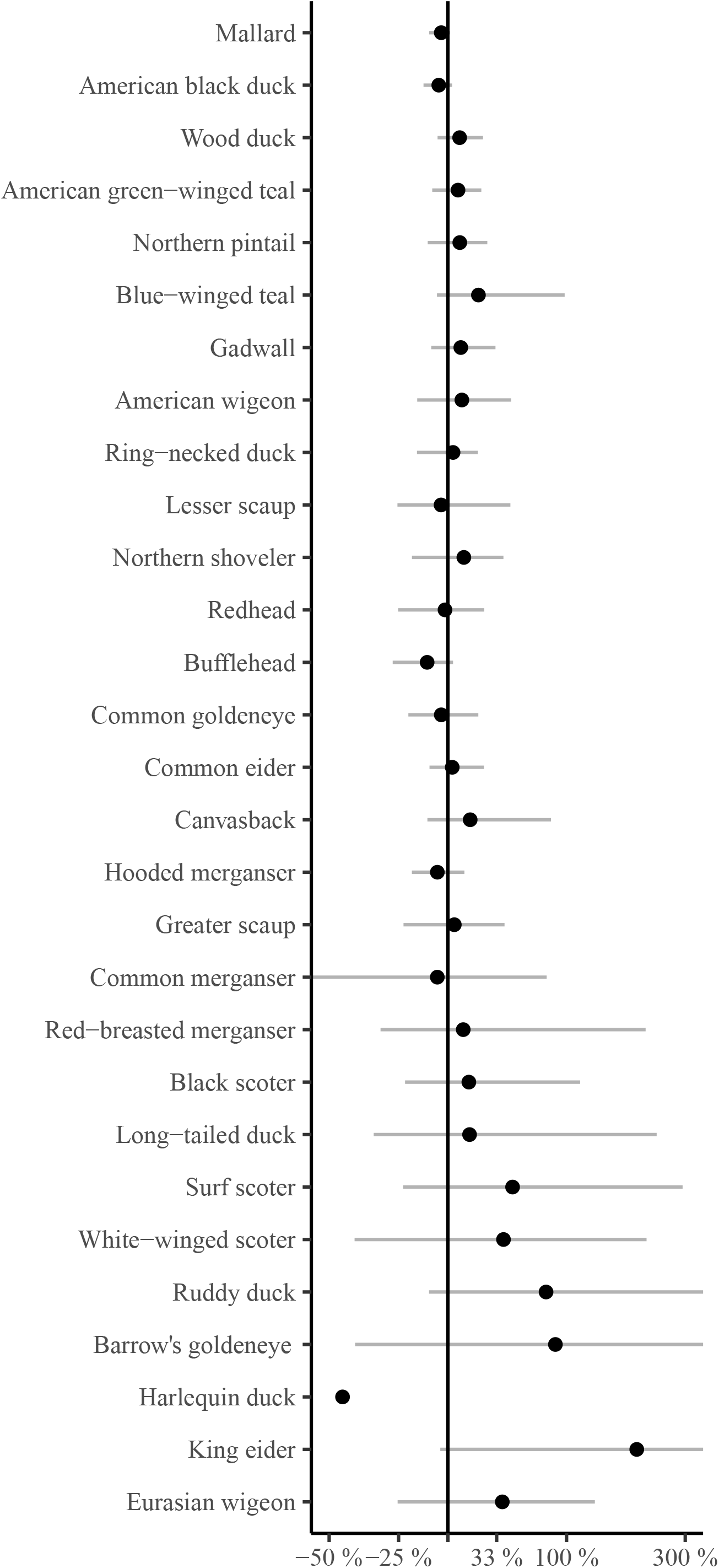
Magnitude of the national harvest estimate from the new model as a percentage of the old model estimate, for ducks from 2010-2019. Positive values indicate new model estimated greater harvest than the old model, negative values are the reverse. Points represent the geometric means and the error bars show the range over the ten-years. Species are sorted based on their total national harvest from the most commonly harvested species at the top (mallard [*Anas platyrhynchos*] and American black duck [*Anas rubripes*]), to the least commonly harvested species at the bottom (e.g., harlequin duck [*Histrionicus histrionicus*], king eider [*Somateria spectabilis*], and Eurasian wigeon [*Mareca penelope*]). For some species near the bottom of the plot, the plotted values only include years in which the old model estimated some harvest of the species (to avoid infinite ratios), including harlequin duck where the old model only estimated some non-zero harvest in one year. Other species on the plot include, wood duck [*Aix sponsa*], American green-winged teal [*Anas carolinensis*], northern pintail [*Anas acuta*], blue-winged teal [*Anas discors*], gadwall [*Mareca strepera*], American wigeon [*Mareca americana*], ring-necked duck [*Aythya collaris*], lesser scaup [*Aythya affinis*], northern shoveler [*Spatula clypeata*], redhead [*Aythya americana*], bufflehead [*Bucephala albeola*], common goldeneye [*Bucephala clangula*], common eider [*Somateria mollissima*], canvasback [*Aythya valisineria*], hooded merganser [*Lophodytes cucullatus*], greater scaup [*Aythya marila*], common merganser [*Mergus merganser*], red-breasted merganser [*Mergus serrator*], black scoter [*Melanitta americana*], long-tailed duck [*Clangula hyemalis*], surf scoter [*Melanitta perspicillata*], white-winged scoter [*Melanitta deglandi*], ruddy duck [*Oxyura jamaicensis*], and Barrow’s goldeneye [*Bucephala islandica*].

**Figure 4.**
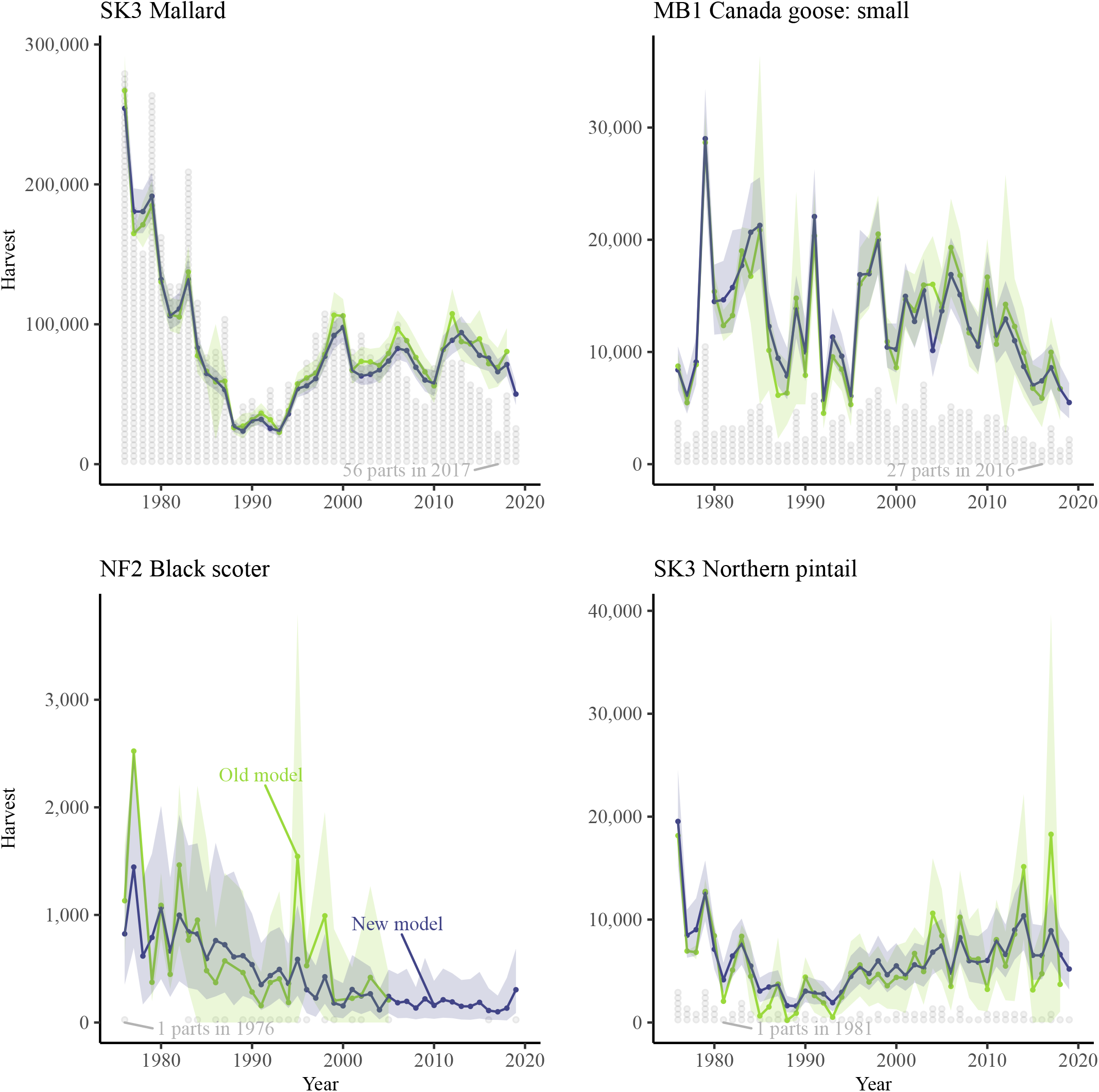
Examples of zone-level estimates of the species-specific harvest for a selection of waterfowl species that range from relatively data-rich (mallard and Canada goose: small [mostly *Branta hutchinsii* ^a^]) to relatively data-poor (black scoter and northern pintail). Estimates are included for all years from 1976-2019 from the Canadian National Harvest Survey, using the new hierarchical Bayesian model described here (darker line) and the old model that it replaces (lighter line). The semi-transparent ribbon surrounding each line represents the 95% credible/confidence interval on the estimates. The light-grey stacked dots represent the number of individual parts for each species submitted in each year (each dot represents 10 wings or tail-fans), and the grey labels indicate the lowest non-zero count of parts for a given species. Province and zone abbreviations are SK3 = Saskatchewan zone 3, MB1 = Manitoba zone 1, and NF2 = Newfoundland and Labrador zone 2. ^a^ This species identification is approximate, because taxonomy and identifications from submitted parts have changed over the course of the survey.

**Figure 5.**
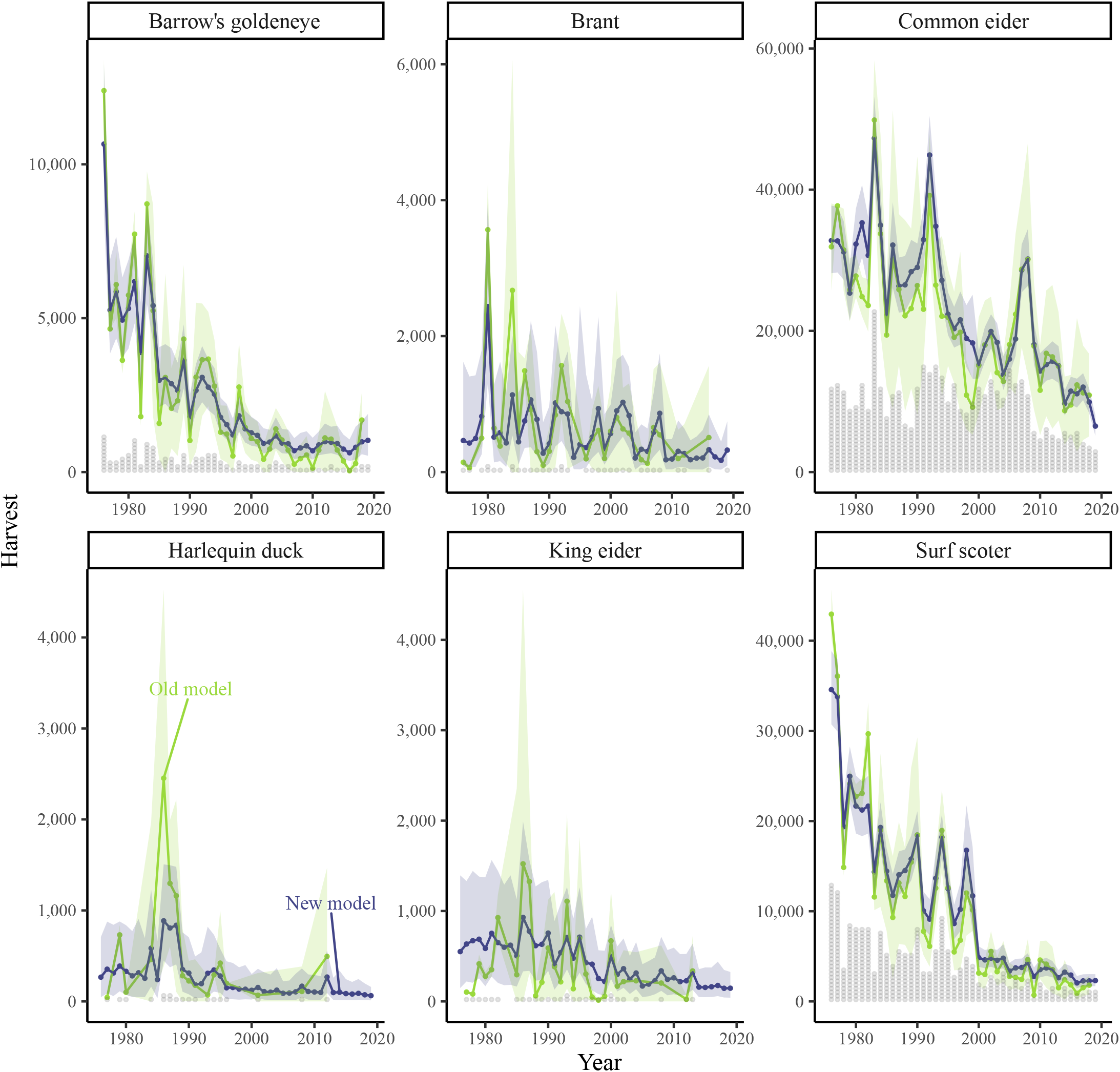
Examples of national estimates of the species-specific harvest for a selection of species that are relatively data-poor and some are of conservation concern (e.g., harlequin duck and Barrow’s goldeneye). The new model generates estimates for all species in all years and they fluctuate less among years. Estimates are included for all years from 1976-2019 from the Canadian National Harvest Survey, using the new hierarchical Bayesian model described here (darker line) and the old model that it replaces (lighter line). The semi-transparent ribbon surrounding each line represents the 95% credible/confidence interval on the estimates. The light-grey stacked dots represent the number of individual parts for each species submitted in each year (each dot represents approximately 10 wings or tail-fans).

The new model estimates are generally more precise than the old model estimates, both for the data-rich species (e.g., mallard and American black duck; Fig.6) and for the relatively data-sparse species (e.g., common eider and black scoter; Fig.6). Averaged across all species of ducks and geese, the coefficients of variation for national harvest estimates are reduced by approximately 10% with the new model. With the new model, 23 species of ducks and geese have average annual national harvest estimates with coefficients of variation less than 10% over the last decade (2010-2019), in comparison with only 6 species using the old model. In addition, the precision of the new model estimates is much more stable through time (i.e., smaller annual fluctuations) and is more precise, relative to the old model, in recent years, when sample sizes are much lower than they were at the beginning of the survey.

**Figure 6.**
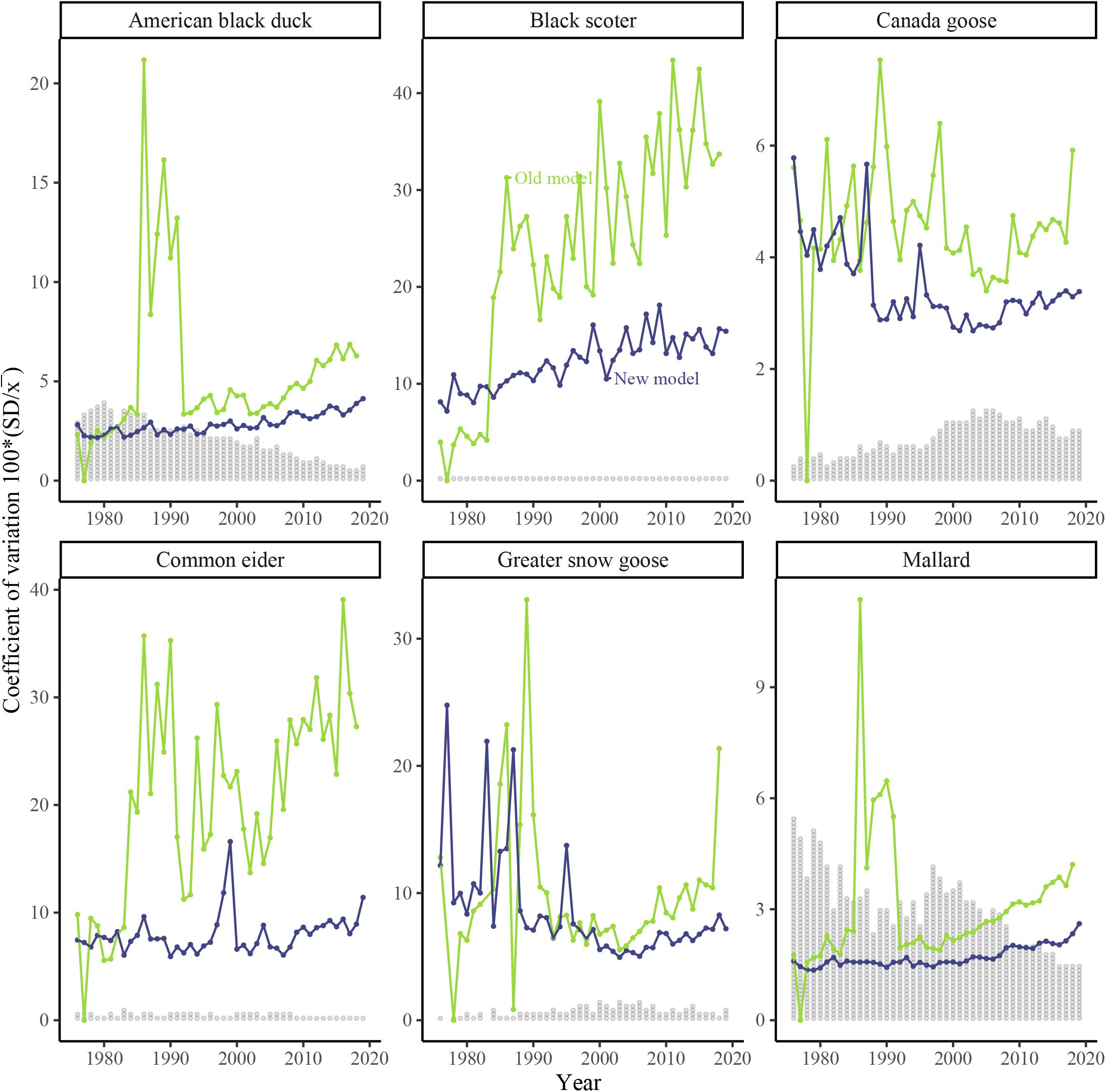
Uncertainty in national estimates of the species-specific harvest for a selection of species that are both relatively precise (e.g., mallard and American black duck) and relatively imprecise (e.g., black scoter and common eider). Estimates are included for all years from 1976-2019 from the Canadian National Harvest Survey, using the new hierarchical Bayesian model described here (darker line) and the old model that it replaces (lighter line). The CV values for the new model are lower (higher precision) in most years and the difference between the two models is increasing through time. Note: the large fluctuations in the old model results in late 1980s and early 1990s reflect annual variations in sampling rates across hunter classes and geographic strata. The light-grey stacked dots represent the number of individual parts for each species submitted in each year (each dot represents up to 200 wings or tail-fans).

With the new model, we also generate formal estimates of age ratios in the harvest, which have estimates of uncertainty and are similarly less sensitive to annual fluctuations (Fig. 7), compared to the raw summaries of parts received from the old model. For example, national-scale age ratios for mallards from the new model are all lower than ratios from the old model (Fig. 8). The new model more accurately weights the contribution of parts from different regions. For example, Ontario and Saskatchewan vary a great deal in both the observed age ratios in the parts and the participation rates in the SCS. Specifically, in Ontario, survey participants have traditionally submitted almost three times as many parts in relation to the total harvest, than have been submitted in western provinces such as Saskatchewan. The new model accounts for the bias in participation rates and generates unbiased estimates of the national age ratios (Fig. 8).

**Figure 7.**
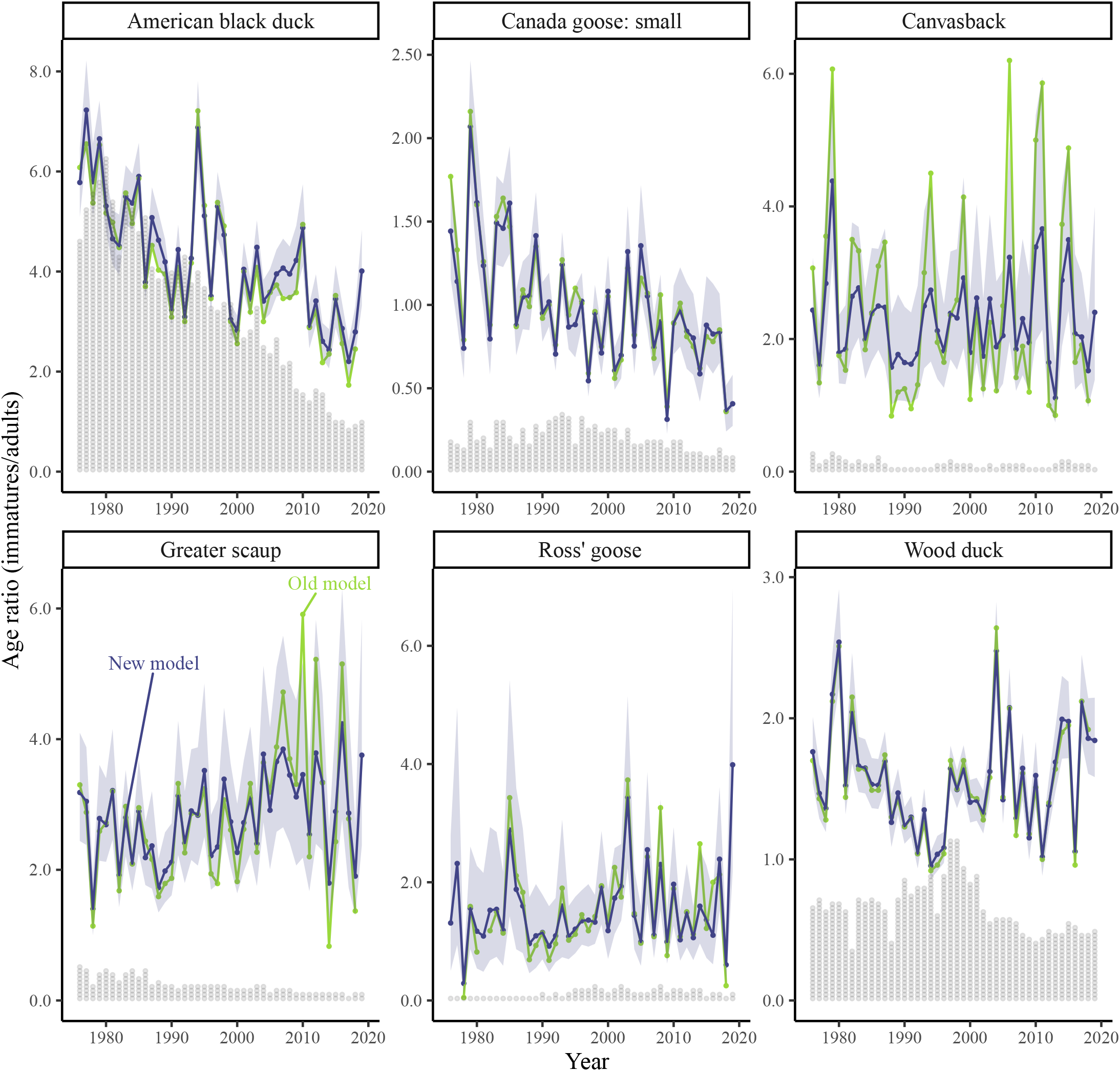
Examples of national estimates of the species-specific age ratios in the harvest for a selection of species that are relatively data-rich (American black duck, Canada goose: small and wood duck) and relatively data-poor (canvasback, greater scaup, and Ross’s goose [*Chen rossii*]). The new model generates estimates of age ratios that are less sensitive to sampling error among years and include estimates of uncertainty. Estimates are included for all years from 1976-2019 from the Canadian National Harvest Survey, using the new hierarchical Bayesian model described here (darker line) and the old model that it replaces (lighter line). The semi-transparent ribbon surrounding each line represents the 95% credible interval on the estimates. The light-grey stacked dots represent the number of individual parts for each species submitted in each year (each dot represents approximately 50 wings or tail-fans). ^a^ This species identification is approximate, because taxonomy and identifications from submitted parts have changed over the course of the survey.

**Figure 8.**
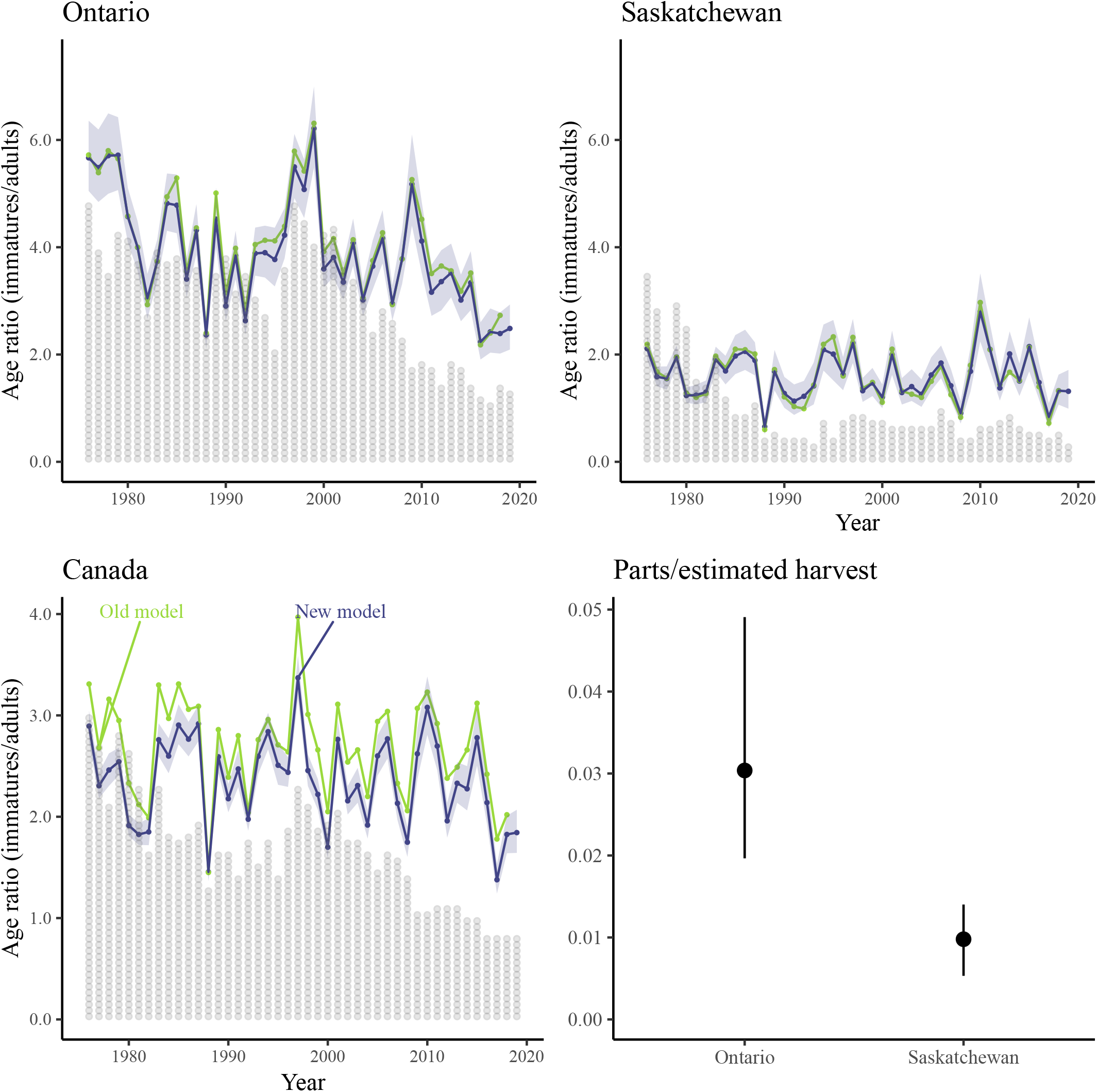
Example of the reduced bias in estimates of national age ratios because the new model adjusts the national estimates for the harvests and number of parts submitted among hunting zones. In this example, the national age ratios for mallard from the old model were biased by the greater number of parts submitted / bird harvested (parts / harvest) and the higher age ratios that tend to occur in eastern Canada (e.g., Ontario) as compared to western Canada (e.g., Saskatchewan). The new model adjusts the national estimates for the relative harvest and parts submission rates, and so removes this bias. Estimates are included for all years from 1976-2019 from the Canadian National Harvest Survey, using the new hierarchical Bayesian model described here (darker line) and the old model that it replaces (lighter line). The semi-transparent ribbon surrounding each line represents the 95% credible interval on the estimates. The light-grey stacked dots represent the number of individual parts for each species submitted in each year (each dot represents 200 wings).

## DISCUSSION

This new model represents an important advance in estimating and managing the harvest of migratory birds. The general benefits of a hierarchical Bayesian framework are well recognized in ecological data analysis (Cressie et al. 2009, Link and Barker 2010, Gelman et al. 2013, Dorazio 2016), and recently in estimating harvest (Arnold 2019), or harvest rate using American black duck band returns (Conroy et al. 2005, Roy et al. 2015). By assuming that many of the key components of the harvest in a given year are similar to that same component in the previous year, the model allows annual estimates to vary through time, but dampens large annual fluctuations due to sampling noise. As a result, the model generates estimates of migratory bird harvest that are less susceptible to sampling error in a given year, particularly for species that occur less frequently in the species composition survey (e.g., Koneff et al. 2017). These less common species include some that have particularly important conservation and management concerns, including harlequin duck (*Histrionicus histrionicus*) and Barrow’s goldeneye (*Bucephala islandica*): two duck species with eastern populations assessed as “special concern” by the Committee on Status of Endangered Wildlife in Canada (COSEWIC 2001 and COSEWIC 2013), but that still appear sporadically in the harvest. By contrast, the traditional analysis would estimate no harvest for less common species if no parts were submitted in a given year, such as all national estimates for harlequin duck since 2012 (Fig. 5). Harvest estimates from the new model are essentially the same for the most commonly harvested species as estimates from the old model that used a series of design-based equations. The new model estimates are generally more precise, which likely derives from the time-series components of the model that share information through time and take advantage of the similarities from one year to the next in the pool of hunters, regional species composition, hunting preferences and behaviours, and regulation packages. The similarity between the new and old model estimates suggests that there is no need to revisit past management decisions, and that these new estimates can integrate seamlessly with existing decision processes for harvest management.

This new model also provides more realistic and practical estimates of uncertainty than the traditional approach. It includes a coherent data-generating model based on the non-negative counts of harvested birds, days spent hunting, etc., and a log-link component that ensures all predictions of harvest are non-negative. The old model used ratio estimators and their standard errors, and so confidence intervals could not be directly estimated (e.g., a naïve interval would include negative numbers of harvested birds for highly uncertain or low-magnitude estimates). The Bayesian framework ensures that all estimates relate to the posterior distributions of the parameters that are directly of interest (i.e., number of harvested birds) and so uncertainty intervals can be interpreted as a range of values that has a high probability of containing the true value (Gelman et al. 2013, Dorazio 2016). In addition, the new model provides a more comprehensive integration of the various sources of uncertainty, because it has explicitly defined distributions for the data and for all parameters and it propagates all uncertainty in those parameters through to all estimates of harvest, age-ratios, species compositions, etc. (Cressie et al. 2009, Gelman et al. 2013). For example, the addition of age and sex specific harvest estimates and associated variation as a standard model output should be a useful feature for integrating these improved estimates, which were previously unavailable, into subsequent analyses such as Lincoln estimates of population sizes (Alisauskas et al. 2014, Arnold 2019).

We see this new model as an initial step in an ongoing evolution of our estimation and understanding of migratory bird harvest in Canada. The hierarchical Bayesian framework provides a clear and flexible way to customise the model, modify the prior assumptions, and add additional sources of information. We suggest it would be useful to explore for potential bias in the estimates of sex-ratios or the distribution of harvest across the season that may be introduced by assuming that missing data are missing at random. Also, small modifications to the priors and distributional structures of this model would allow for stronger assumptions about species composition across years and between adjacent periods within years. Currently, the priors on the time-series components are only weakly informative (Gelman 2008, van de Schoot et al. 2021), but more informative priors are likely warranted, given the deep domain-specific knowledge that exists in the waterfowl harvest management community (Banner et al. 2020). For example, it may be beneficial to include informative priors that share information among selected species based on knowledge about species exposure to similar hunting behaviour, such as informing the harvest of Harlequin duck using information about the harvest of Common eider, or scoter species. This model framework also makes it relatively easy to add informative covariates that might influence hunting behaviour, such as weather conditions during peak hunting periods. Additionally, the spatial stratification provided by the hunting zones is a useful simplification that captures a great deal of the variation in harvests (Cooch et al. 1978). However, an inherently spatial treatment of the data could allow this model to track the spatial variation while also sharing some information among neighbouring regions (e.g. Morris et al. 2019) and further improve the quality of the estimates in some data-sparse regions. For example, murres are harvested in zones 1 and 2 of the province of Newfoundland and Labrador, but currently we are unable to estimate murre harvest for zone 2, because there are not enough parts or survey responses. Efforts are underway to increase response rates for the survey and to generate more data to help fill this and other gaps. With these additional data and by sharing information in a hierarchical framework and assuming that some aspects of the murre harvest are similar between zone 2 and zone 1 of the province, we can further improve our estimates.

One of the main areas for future improvements would be a better understanding of the variation among hunters in the species composition of their harvest. Currently, both the new model and the old model pool all parts contributed in a given zone and period, regardless of which hunter contributed the parts. This simplification has been necessary because of the relatively low participation rate of hunters in the SCS (Cooch et al. 1978), the relatively large number of parts required to estimate species proportions in each period of the hunting season, and the complexities of coherently tracking uncertainties in the old model. However, the new model’s hierarchical Bayesian framework provides a clear way to account for this additional source of variation. Future research could consider adding a sub-model to estimate and account for among-hunter variation. This sub-model could share information among years, seasons, and zones. It could also benefit from potential modifications to the surveys to collect additional information on hunter behaviour, such as identifying hunters that specialize on hunting seaducks or that regularly hunt in the winter. In addition, with future plans to transition the survey instrument from its current hardcopy format to an electronic format, it is hoped that individual hunters can be tracked through time as multiple years of information on each hunter would greatly improve our ability to account for the variation among hunters. Similarly, improved accounting for variations among hunters and this new modeling approach provide a coherent framework to integrate estimates of possible response bias and the uncertainty in those estimates of bias (Padding and Royle 2012).

## MANAGEMENT IMPLICATIONS

With this new model and modeling framework, we have generated improved estimates of migratory game bird harvest, particularly for species that are less commonly harvested. The hierarchical nature of the new model and the time-series components that allow annual variation while sharing information through time make more efficient use of the survey data than the old model that treated each year as independent. The new model also makes it possible to estimate age and sex specific harvests that will facilitate more informed management decisions (e.g., Alisauskas et al. 2014). The Bayesian model propagates uncertainty across all the parameters in a coherent and transparent way, so that management decisions can more fully integrate that uncertainty (Nichols et al. 1995). The Bayesian nature of the new model, and the open-access code in the supplement, provides a coherent framework to incorporate the rich prior knowledge of hunting behaviour and species biology, which will facilitate future improvements and elaborations to fill more specific management information needs (Banner et al. 2020).

*Associate Editor*

## Summary for online Table of Contents

This new model provides improved estimates of the hunting activity and harvest of migratory game birds in Canada, as well as new estimates for age- and sex-specific harvest. The Bayesian framework and the open-source code that we provide here will facilitate ongoing improvements and elaborations of the model.

## LITERATURE CITED

Ahlers, A., and C. Miller. 2019. Testing waterfowl hunters’ waterfowl identification skills. Wildfowl 69: 206-220. Alisauskas, R. T., T.W. Arnold, J. O. Leafloor, D. L. Otis, and J. S. Sedinger. 2014. Lincoln estimates of mallard (Anas platyrhynchos) abundance in North America. Ecology and Evolution 4: 132–143.

Alisauskas, R. T., K. L. Drake, and J. D. Nichols. 2009. Filling a Void: Abundance Estimation of North American Populations of Arctic Geese Using Hunter Recoveries. Pages 463–489 in D. L. Thompson, E. G. Cooch, M. L. Conroy, editors. Modeling Demographic Processes in Marked Populations. Environmental and Ecological Statistics. Arnold, T. W. 2019. A bayesian hierarchical model for estimating American woodcock harvest. Pages 27–34 in D.G. Krementz, D.E. Andersen, and T.R. Cooper, Editors. Proceedings of the Eleventh American Woodcock Symposium. University of Minnesota Libraries Publishing, Minneapolis.

Banner, K. M., K.M. Irvine, and T. J. Rodhouse. 2020. The use of Bayesian priors in ecology: The good, the bad and the not great. Methods in Ecology and Evolution 11: 882–889.

Canadian Wildlife Service Waterfowl Committee. 2020. Population status of migratory game birds in Canada: November 2019. CWS Migratory Birds Regulatory Report Number 52.

Carney, S.M. 1992. Species, age, and sex identification of ducks using wing plumage. U.S. Fish and Wildlife Service, Washington, D.C.

Conn, P.B., D.S. Johnson, P. J. Williams, S. R. Melin, M. B. Hooten, 2018. A guide to Bayesian model checking for ecologists. Ecological Monographs 88: 526–542.

Conroy, M. J., C. J. Fonnesbeck, N. L. Zimpfer. 2005. Modeling regional waterfowl harvest rates using Markov chain Monte Carlo. The Journal of Wildlife Management, 69:77–90.

Cooch, F.G., S. Wendt, G.E.J. Smith, G., Butler. 1978. The Canada migratory game bird hunting permit and associated surveys. Pages 8–41 in H. Boyd and G. Finney, eds. Migratory game bird hunters and hunting in Canada. Canadian Wildlife Service. Occasional Papers. No. 43.

COSEWIC. 2000. COSEWIC assessment and status report on the Barrow’s goldeneye Bucephala islandica, eastern population, in Canada. Committee on the Status of Endangered Wildlife in Canada. Ottawa.

COSEWIC. 2013. COSEWIC assessment and status report on the harlequin duck Histrionicus histrionicus Eastern population in Canada. Committee on the Status of Endangered Wildlife in Canada. Ottawa.

Cressie, N., C.A. Calder, J.S. Clark, J.M.V. Hoef, and C.K. Wikle. 2009. Accounting for uncertainty in ecological analysis: The strengths and limitations of hierarchical statistical modeling. Ecological Applications 19: 553–570.

Dorazio, R. M. 2016. Bayesian data analysis in population ecology: Motivations, methods, and benefits. Population Ecology 58: 31–44.

Gilliland, S.G., H.G. Gilchrist, R.F. Rockwell, G.J. Robertson, J-P.L. Savard, F. Merkel, and A. Mosbech. 2009. Evaluating the sustainability of harvest among northern common eiders Somateria mollissima borealis in Greenland and Canada. Wildlife Biology 15: 24–36.

Hagen, C. A., J. E. Sedinger, and C. E. Braun. 2018. Estimating sex-ratio, survival and harvest susceptibility in greater sage-grouse: making the most of hunter harvests. Wildlife Biology 2018: wlb.00362

Kellner, K. 2019. jagsUI: A Wrapper Around ‘rjags’ to Streamline ‘JAGS’ Analyses. R package version 1.5.1. https://CRAN.R-project.org/package=jagsUI

Osnas, E. E., Q. Zhao, M. C. Runge, and G. S. Boomer (2016). Cross-seasonal effects on waterfowl productivity: Implications under climate change. Journal of Wildlife Management 80:1227–1241.

Link, W.A., J.R. Sauer, D.K. Niven. 2020. Model selection for the North American Breeding Bird Survey. Ecological Applications, 30:e02137.

Koneff, M. D., G. S. Zimmerman, C. P. Dwyer, K. K. Fleming, P. I. Padding, P. K. Devers, F. A. Johnson, M. C. Runge, and A. J. Roberts. 2017. Evaluation of harvest and information needs for North American sea ducks. PLOS ONE 12: e0175411.

Morris, M., K. Wheeler-Martin, D. Simpson, S. J. Mooney, A. Gelman, and C. DiMaggio. 2019. Bayesian hierarchical spatial models: Implementing the Besag York Mollié model in stan. Spatial and Spatio-Temporal Epidemiology 31: 100301.

Nichols, J. D., F. A. Johnson, and B. K. Williams. 1995. Managing North American waterfowl in the face of uncertainty. Annual Review of Ecological Systems 26:177–99

North American Waterfowl Management Plan Committee NAWMP. 2018. North American waterfowl management plan: Connecting people, waterfowl, and wetlands. U.S. Department of the Interior, Environment and Climate Change Canada, and Environment and Natural Resources Mexico, Department of the Interior, Washington, D.C., USA.

Padding, P. I., and J. A. Royle. 2012. Assessment of bias in US waterfowl harvest estimates. Wildlife Research 39: 336–342.

Palumbo, M.D., J.W. Kusack, D.C. Tozer, S.W. Meyer, C. Roy, and K.A. Hobson. 2020. Source areas of blue-winged teal harvested in Ontario and Prairie Canada based on stable isotopes: implications for sustainable management. Journal of Field Ornithology 91: 64–76.

Plummer, M. 2003. JAGS: A program for analysis of Bayesian graphical models using Gibbs sampling. Proceedings of the Third International Workshop on Distributed Statistical Computing, 20-22 March 2003, Vienna, Austria.

Roy, C., S.G. Cumming, E.J.B. McIntire. 2015. Spatial and temporal variation in harvest probabilities for American black duck. Ecology and Evolution, 5:1992–2004.

Saunders, S. P., M.T. Farr, A.D. Wright, C.A. Bahlai, J.W. Ribeiro Jr., S. Rossman, A.L. Sussman, T.W. Arnold, and E.F. Zipkin. 2019. Disentangling data discrepancies with integrated population models. Ecology 100(6):e02714. 10.1002/ecy.2714

Sen, A. R. 1976. Developments in migratory game bird surveys. Journal of the American Statistical Association 71: 43–48.

Sen, A. R., S. Sellers, and G. E. J. Smith. 1975. The use of a ratio estimate in successive sampling. Biometrics 31: 673–683.

Smith, G. E. J. 1974. A study of two methods to increase the efficiency of the species composition survey. Canadian Wildlife Service Biometric section. Manuscript Report No. 11.

van de Schoot, R. S, Depaoli, R. King, B. Kramer, K. Märtens, M.G. Tadesse, M. Vannucci, A. Gelman, D. Veen, J. Willemsen, and C. Yau. 2021. Bayesian statistics and modelling. Nature Reviews Methods Primers 1: 1–26.

Zimmerman, G.S., W.A. Link, M.J. Conroy, J.R. Sauer, K.D. Richkus, and G.S. Boomer. 2010. Estimating migratory game-bird productivity by integrating age ratio and banding data. Wildlife Research 37:612–622.

Zimmerman, G.S., J.R Sauer, G.S. Boomer, P.K. Devers, and P.R. Garrettson. 2017. Integrating Breeding Bird Survey and demographic data to estimate Wood Duck population size in the Atlantic Flyway. The Condor 119: 616–628

Zipkin, E. F., E. R. Zylstra, A. D. Wright, S. P. Saunders, A. O. Finley, M. C. Dietze, M. S. Itter, M. W. Tingley. 2021. Addressing data integration challenges to link ecological processes across scales. Frontiers in Ecology and the Environment, 19:30–38.

